# Activity Budgets and Boldness as Potential Predictors of Release Readiness in Rehabilitating Javan Slow Lorises

**DOI:** 10.64898/2025.12.28.696733

**Authors:** Abdullah Langgeng, Marie Sigaud, Wendi Prameswari, Nur Purba Priambada, Puji Rianti, Richard Moore, Ikki Matsuda, Andrew MacIntosh

## Abstract

Rehabilitation centers are central to the conservation of Javan slow lorises (*Nycticebus javanicus*) by providing care for individuals confiscated from trade and preparing them for potential release. To evaluate behavioral readiness for reintroduction, we quantified activity budgets and boldness in 10 adult *N. javanicus* (7 females and 3 males) housed at a rehabilitation facility in West Java, Indonesia, from May to September 2024. Locomotion dominated activity budgets (57.14% ± 24.86%), followed by resting (23.51% ± 16.61%), with low rates of stereotypy (4.36% ± 13.48%) and grooming (2.33% ± 5.28%). Boldness tests using six novel object types revealed strong stimulus dependence: predator models suppressed approach, while enrichment and neutral objects encouraged exploration. Sex, housing conditions, and rehabilitation duration did not predict boldness, indicating inter-individual differences in responses to novelty. Principal component analysis identified an integrated bold–exploratory activity axis, suggesting that activity budgets and boldness covary as part of a behavioral syndrome rather than independent traits. Together, these results indicate that combined activity and personality metrics provide complementary indicators of behavioral welfare and release readiness, with potential to improve pre-release assessment and site matching for this critically endangered primate.

## Introduction

Across Southeast Asia, the growing illegal wildlife trade and widespread habitat loss have created an urgent need for wildlife rescue and rehabilitation centers (Cheyne 2006; Nijman 2009). The region is recognized as one of the world’s major wildlife trade hotspots, with millions of animals taken from the wild each year for the exotic pet and traditional medicine markets (Nijman 2009). In Indonesia alone, slow lorises, gibbons, macaques, and many bird species are frequently confiscated by authorities from traders or private owners. Consequently, wildlife rescue centers have become indispensable institutions that provide immediate care and long-term rehabilitation for displaced or confiscated animals (Moore et al. 2014). These facilities function both as sanctuaries and as transitional spaces, bridging animal welfare and conservation by preparing individuals for reintroduction once their physical and behavioral conditions permit (Cheyne 2006; EAZA 2022). Beyond individual welfare, rescue centers support conservation at a broader scale through law enforcement, public education, and the development of scientifically informed reintroduction protocols. The International Union for Conservation of Nature (IUCN) recognizes well-managed translocations as valuable conservation tools when accompanied by thorough health, behavioral, and habitat assessments that ensure welfare and post-release survival (IUCN 2013). Consequently, wildlife rescue centers have evolved from reactive welfare facilities into key components of integrated species recovery strategies across Southeast Asia (EAZA 2022; Moore et al. 2014).

Despite their importance, the success of reintroduction programs remains inconsistent among species and regions. Post-release mortality in primates and other mammals is often high, commonly resulting from behavioral maladaptation, stress, or inadequate preparation for wild conditions (Kenyon et al. 2014; van der Sandt 2017). While veterinary screening, including checks for injuries, parasites, or disease, is a standard part of pre-release procedures, behavioral readiness has received comparatively little systematic attention (IUCN-SSC 2019). Activity budgets, which quantify how animals allocate time among key behaviors, are practical tools for assessing behavioral welfare and environmental suitability in captivity and for evaluating the readiness of rehabilitated individuals for release through comparison with wild counterparts (Maple and Perdue 2013; Melfi and Feistner 2002). Deviations from species-typical activity patterns, including excessive resting or stereotypic behaviors, may indicate environmental inadequacy, whereas balanced expression of natural behaviors suggests effective enrichment and enclosure design (Maple and Perdue 2013; Melfi and Feistner 2002). Thus, activity-budget analysis is useful both for evaluating the quality of captive environments and for predicting behavioral readiness for reintroduction.

In addition to activity-based measures, individual behavioral tendencies such as boldness and exploration also influence post-release adjustment and therefore complement activity budgets as indicators of behavioral readiness for release (Bremner-Harrison et al. 2004; Reale et al. 2007; Wolf and Weissing 2012). Personality, consistent individual differences in behavior across time and context (Dingemanse et al. 2010; Gosling 2001; Morita et al. 2020; Sih et al. 2004a), has been repeatedly linked to fitness and translocation outcomes (de Azevedo and Young 2021; Haage et al. 2017). Bold individuals may explore new environments more readily but can also suffer higher predation risk or mortality, whereas shy individuals are more cautious but may obtain fewer resources (Bremner-Harrison et al. 2004; Carter et al. 2013; Haage et al. 2017). The adaptive value of such traits possibly depends on ecological conditions (de Azevedo and Young 2021). Incorporating personality assessment into pre-release evaluations would improve the selection of individuals whose behavioral tendencies match the ecological and social conditions of specific release sites (Carter et al. 2013; Langridge et al. 2021; Wolf and Weissing 2012).

Beyond examining boldness in isolation, understanding whether behavioral traits covary, forming behavioral syndromes, provides deeper insight into individual coping strategies and release success potential. Behavioral syndromes, defined as suites of correlated behaviors expressed consistently across contexts, reflect integrated personality profiles that shape how individuals navigate environmental challenges (Sih et al. 2004b). For instance, bold-exploratory syndromes may link high locomotor activity with increased novel object investigation, whereas shy-cautious syndromes may couple reduced exploration with prolonged vigilance (Fior et al. 2018; Kudo et al. 2021). Identifying such syndromes in rehabilitating primates could reveal whether activity budgets and boldness responses reflect coherent personality dimensions, thereby improving predictions of post-release performance and informing individualized management strategies.

Slow lorises (*Nycticebus* and *Xanthonycticebus*: Nekaris and Nijman 2022) are small nocturnal primates endemic to South and Southeast Asia, distinguished by their cryptic movements, strong grasping ability, and venomous bite (Nekaris and Burrows 2020). They are largely solitary or live in loosely connected social networks characterized by mutual avoidance and occasional affiliative interactions (Nekaris and Bearder 2011). Slow lorises face severe threats from habitat loss, poaching, and the illegal pet trade, with most species now listed as Endangered or Critically Endangered (IUCN 2025; Nekaris and Starr 2015). The Javan slow loris (*N. javanicus*) is among the slow loris species frequently observed in wildlife markets and confiscations in Indonesia (Nekaris and Jaffe 2007). Rehabilitation programs, therefore, are central to conservation efforts for this species (EAZA 2022; Moore et al. 2014). However, despite increased attention and substantial investment, the success of reintroduction programs for slow lorises remains limited, with high mortality rates and inconsistent behavioral outcomes (Moore et al. 2014; Streicher et al. 2003; van der Sandt 2017).

Recent work shows that pygmy slow lorises (*X. pygmaeus*) exhibit more affiliative and socially flexible behavior than once assumed, with social housing improving welfare for both sexes (Yamanashi et al. 2021). These findings challenge the long-standing view of slow lorises as strictly solitary and highlight the importance of behavior-based rehabilitation. Although slow loris rehabilitation has expanded, successful reintroductions remain limited due to insufficient behavioral preparation, habitat mismatch at release sites, and a lack of standardized pre-release assessment (EAZA 2022; Moore et al. 2014). Naturalistic diets, increased enclosure complexity and nocturnal behavioral monitoring have been shown to improve welfare and promote more natural activity patterns in captive slow lorises (Al-Razi et al. 2020; Alejandro et al. 2021; Williams et al. 2015), yet activity budgets and personality metrics are rarely combined to evaluate release readiness. As shown in other primate species such as gibbons, reintroduction success depends on behavioral competence as well as habitat suitability (Cheyne 2006), and the cryptic nocturnal habits of slow lorises make such assessment especially crucial.

Here, we evaluate the behavioral readiness of confiscated *N. javanicus* at rescue center in West Java, Indonesia, by quantifying activity budgets and boldness traits using novel-object tests. Specifically, we examine whether behavioral traits covary to form personality syndromes and assess how intrinsic personality dimensions and extrinsic factors (housing, enclosure characteristics) shape behavioral expression. By integrating activity budgets and boldness into a unified personality framework, we aim to identify behavioral indicators relevant to welfare and post-release performance, contributing to more effective rehabilitation and conservation of this critically endangered primate.

## Methods

### Study site and subjects

The study was conducted at Yayasan Inisiasi Alam Rehabilitasi Indonesia (YIARI), a slow loris rescue center located in Bogor, West Java, Indonesia, which houses over 100 confiscated *Nycticebus* spp. The research involved ten adults of *N. javanicus*, consisting of seven females and three males, selected as study subjects and assigned with a unique identifier (i.e. JS01–JS10). These individuals were deemed for translocation within the subsequent months. The subjects were housed under varying conditions, including differences in enclosure size, presence of conspecifics, and length of stay at the rehabilitation center (ranging from approximately 4 to 32 months at study onset; see supplemental Table S1 for details of housing conditions and stay length). The *N. javanicus* at YIARI were provided with daily provisions of vegetables, fruits, gums, insects, along with enrichment activities. All subjects housed in a group were individually identifiable by unique facial and physical characteristics, including facial markings, scars, and coat coloration.

### Behavioral data collection

#### Activity budget

We conducted behavioral observations at night, during *N. javanicus* active period (17:00–05:00) from May to September 2024. Behavioral observations of each subject were balanced as much as possible across nights and time slots to ensure representative sampling of activity budgets throughout the nights. Data on 10 adult individuals were collected using an instantaneous focal sampling technique in which the behavior of the focal individual was recorded every 1 minute over a 45-minute observation period. We collected the data using a tablet (Samsung Galaxy A9+) installed with ZooMonitor software ((https://zoomonitor.org/; Lincoln Park Zoo). Behaviors were classified on Table 1.

**Table 1.** Ethogram used for activity-budget scoring. Behavioral categories and operational definitions used for instantaneous focal sampling (1-min intervals)

| Behaviours | Definition |
| --- | --- |
| Locomotion | Movement associated with looking for food (often includes visual and olfactory searching), or exploring the enclosure |
| Resting | Remain stationary, often with body hunched, eyes open or sleeping |
| Feeding | Actual consumption of a food item |
| Stereotypy | Repetitive and no obvious goal behaviors indicating distress (e.g. pacing, circling, excessive grooming) |
| Self-grooming | Autogroom, lick or use tooth comb on own fur |
| Social | All interactions with conspecifics, including aggression, allogroom, play and other social behaviours |
| Alert | Remain stationary like in “rest” but active observation of environment or observer |
| Others | Other behavior not included in ethogram |
| Out of Sight | Individual cannot be seen |

#### Behavioral experiments for boldness assessments

To assess individual boldness, each subject was exposed to six novel objects representing four different types of stimuli: predator models (snake and leopard cat plush toys), positive stimuli (two novel enrichments: bottled honey and a fruit bamboo puzzle), a conspecific stimulus (mirror), and a neutral stimulus (pyramid object) (supplemental Figure S1). Prior to testing, individuals had no exposure to the test objects or similar items. We exposed each study subject to each object only once, in a randomized order, with the object set up prior to their waking time (16:30 – 17:00). The data collection began once the focal individual emerged from their nesting box and acknowledged the presence of the objects. Behaviors were categorized into four levels of risk-taking: very bold, bold, shy, and very shy, with each category assigned a corresponding score (Table 2: Bremner-Harrison et al. 2018). Using continuous focal sampling, behaviors were recorded over a 15-minute period to calculate a boldness index. This duration was selected to minimize stress in potentially neophobic individuals while allowing sufficient time for initial approach and object investigation, based on pilot observations. To mitigate potential adverse effects of novel-object exposure, a before–after observational approach was used to monitor short-term changes in behavior immediately following object presentation.

**Table 2.** Boldness scoring system. Risk-taking levels assigned to behaviors during novel-object trials. Positive scores denote approach/exploration, negative scores denote withdrawal/avoidance

| **Extremely bold** | **Bold** | **Shy** | **Extremely shy** |
| --- | --- | --- | --- |
| Bold approach (approach within 1 body length) | Approach (normal pace approach >1 body length) | Hesitant approach (approach slowly, with multiple pauses, >1 body length) | Remain stationery & alert (complete stillness for at least 3 seconds, with eyes alert) |
| Investigate object (constantly observe the object within 1 body length) | Rest relaxed (inactive with eyes drooped or closed) | Rest alert (rest with eyes alert) | Venom pose (display brachial gland) |
| Touch object (make brief physical contact with object) | Autogroom (self-clean) | Watch observer (intently observe the observer) | Run away (flee quickly from object) |
| Sniff object (smell the object) | Feeding (ingestion of and /or search food) | Back away (move away slowly in a backward manner from object) | Scream (loud vocalization indicating fear) |
| Play object (make long physical contact with objects or in bouts) | Whistle (loud monosyllabic vocalization as a contact seeking call) | Chitter (short rhythmic clicks, often used in defense) | Stereotypy (repetitive and no obvious goal behaviors indicating distress or fear) |
| Carry object (grab and move with object attached) |  |  |  |
| Attack object (agonistic contact, i.e. grasping, wrestling, biting, or hitting) the object |  |  |  |

extremely bold = 2, bold = 1, shy = -1, extremely shy = -2

We additionally quantified boldness using a spatially explicit metric by recording each individual’s minimum approach distance to the object measured with a calibrated rangefinder (iOS Measure app). Proximity to a novel object is widely employed as an index of risk-taking, complementing behavioral assessments by indicating how closely an animal would approach a potential threat or unfamiliar stimulus.

### Data analysis

#### Activity budget

To analyze the activity budgets, we calculated the proportion of time each individual spent in the predefined behavioral categories (Table 1). These were derived from focal observations collected during the focal instantaneous sampling. For each individual, the total duration of each behavior was summed and divided by the total observation time to yield a proportional activity budget.

#### Boldness score

Behaviors were scored based on their level of risk-taking: very bold (+2), bold (+1), shy (−1), and very shy (−2) for each test. The duration of each displayed behavior was multiplied by its corresponding score. An individual’s boldness score was then calculated as the average score across the six novel object tests.

#### Statistical analysis

We performed Dirichlet test to evaluate the effects of sex and housing condition on activity budgets using DirichletReg package (Maier 2014). To meet the assumptions of compositional analysis, feeding and social behaviors were pooled into an “other” category. Feeding was scheduled by the rescue center, and social behaviors lacked in solitarily housed individuals, resulting in a high frequency of zero values. Pooling these behaviors reduced zero inflation and allowed more stable estimation of the Dirichlet model. We used Generalized Linear Mixed Models (GLMM) to evaluate the effects of sex, object type, and housing conditions on the boldness index with glmmTMB package (Brooks et al. 2017). To avoid pseudoreplication, individual identity and sampling date were included as random factors in the models. The boldness score for each individual and the closest distance to each object were treated as response variables in boldness and proximity models. Additionally, the relationship between boldness score and the closest distance approached by each individual to the objects was tested using Spearman’s rank correlation coefficient. We used Principal Component Analysis (PCA) to explore patterns of covariation among behavioral measures and to visualize potential behavioral syndromes. Prior to analysis, the suitability of the data for dimension reduction was assessed using the Kaiser–Meyer–Olkin (KMO) measure of sampling adequacy. PCA was conducted on standardized variables using the correlation matrix, and components with eigenvalues greater than 1 were retained for interpretation. Given the small sample size and moderate sampling adequacy, PCA results were interpreted cautiously and treated as exploratory. All statistical analyses were performed in R version 4.4.1 (R-Core-Development-Team 2024). Statistical significance for all analyses was set at p ≤ 0.05.

## Results

### Activity budget

#### Overall

We collected a total of 4,725 data points (78 hours and 45 minutes) across 105 behavioral observations to assess the activity budget of rehabilitating *N. javanicus* (mean ± SD: 7.30 ± 3.21 hours of observations per individual). In general, their activity budget showed that locomotion accounted for the largest proportion of observed behaviors, averaging 57.14% **±** 24.86%. This was followed by resting behaviors, including both resting and sleeping, which together comprised 23.51% ± 16.61% of the activity budget. Other notable behaviors included stereotypic movements (4.36% ± 13.48%), alertness (5.16% ± 6.97%), social interactions (3.32% **±** 7.51%), and feeding (3.13% ± 5.61%). Grooming and other uncategorized behaviors were observed at relatively low frequencies, with averages of 2.33% ± 5.28% and 0.40% ± 3.65% respectively.

#### Males vs. females

Our results indicated variability in the activity budgets, both across different behaviors between sexes and housing conditions (Figure 1). Females spent the largest proportion of time in locomotion (70.11% ± 17.68%) and rest (15.76% ± 11.35%), while stereotypy (0.47% ± 2.51%) and other behaviors (0.79% ± 5.33%) were rare. In contrast, males showed less locomotion (45.70% ± 24.78%) but more rest (30.35% ± 17.58%), with slightly higher feeding and grooming than females. Among males, stereotypy (7.80% ± 17.73%, mostly in solitary individuals) and other behaviors (0.04% ± 0.30%) were the least frequent. A Dirichlet test confirmed that females were significantly more active in locomotion (p = 0.005) than males, but we found no significant difference in other behaviors (p > 0.05).

**Figure 1.**
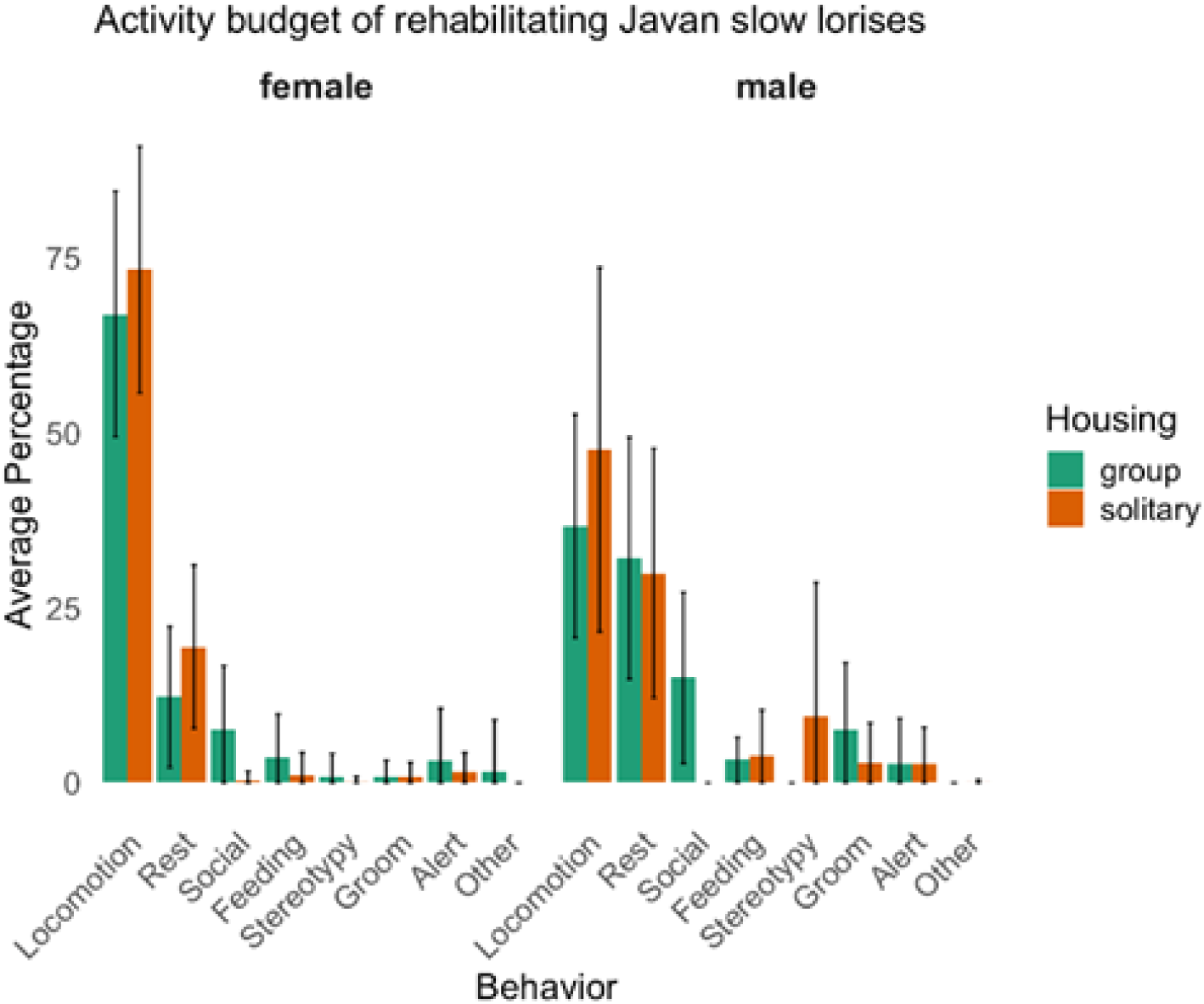
Activity budget of rehabilitating *N. javanicus*. Mean proportion of time allocated to core behaviors during nocturnal observation periods (17:00–05:00). Error bars represent standard deviation across individuals (N = 10).

#### Solitary vs. grouped individuals

Individuals housed solitarily spent the greatest proportion of time in locomotion (56.47% ± 26.38%), with rest as the second most frequent activity (26.35% ± 16.67%). Rare behaviors in this group included grooming (2.16% ± 4.85%) and others (0.03% ± 0.27%), while stereotypy was more common (6.28% ± 16.08%), particularly among solitary males. Grouped individuals likewise showed highest level of locomotion on average (58.48% ± 21.83%), with rest as the second most frequent activity (17.83% ± 15.20%). They also engaged in social interaction (9.77% ± 10.35%). Grooming (2.67% ± 6.13%) and stereotypy (0.52% ± 2.95%) remained infrequent in this group. A Dirichlet test showed no significant effects of housing on the activity budgets of the *N. javanicus* (p > 0.05).

### Boldness

#### Boldness score

A total of 60 experiments (900 minutes in total) were conducted on 10 rehabilitating *N. javanicus* (supplemental Table S2). The results from the GLMM revealed that boldness scores were significantly influenced by the type of object presented. Specifically, *N. javanicus* showed a strong aversion to predator-related objects, exhibiting significantly lower boldness scores. In contrast, subjects exhibited significantly higher boldness scores in association with enrichment objects (Figure 2, Table 3). There was a marginally positive association between boldness score and presentation of the mirror, suggesting some evidence that the appearance of a conspecific (their reflection) induces greater boldness as well. The presence of conspecifics, enclosure size, the duration of stay in the rehabilitation center, and sex were not associated significantly with boldness in this study (Table 3).

**Figure 2.**
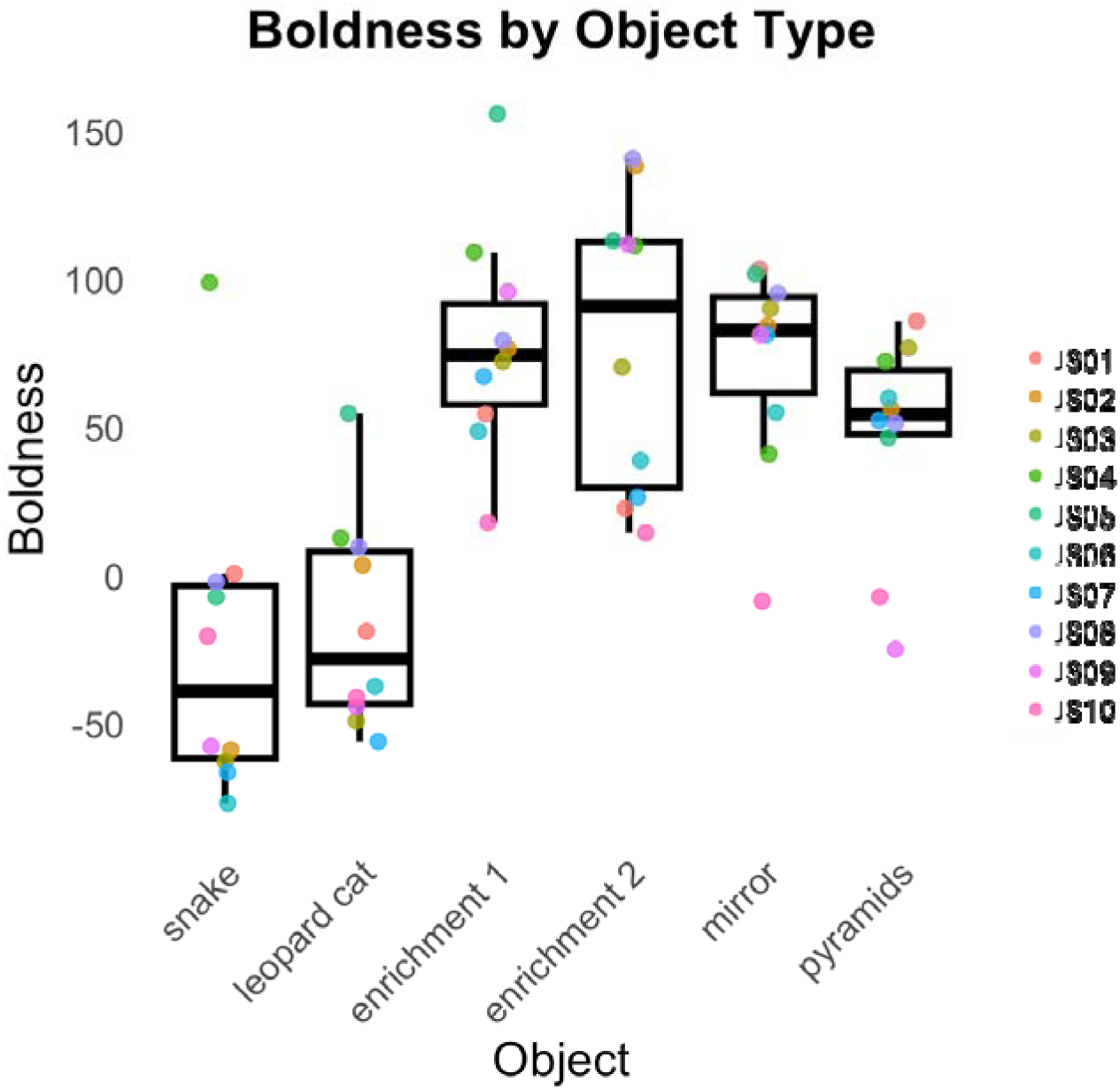
Box plot showing boldness scores of *N. javanicus* (N = 10) presented with six categories of novel objects. Boxes represent the interquartile range between the first and third quartiles, horizontal lines within each box indicate the median, and whiskers show the full range of non-extreme values. Colored points represent individual boldness scores for each object type, illustrating inter-individual variation in responsiveness to novelty.

**Table 3.** Generalized Linear Mixed Model (GLMM) results for predictors of boldness score. Fixed effects included object type, enclosure size, sex, housing type, and length of stay in rehabilitation. Individual identity and sampling date were included as random factors. Significant predictors indicated in bold (p ≤ 0.05).

|  | Estimate | Std. error | z value | Pr(>\|z\|) |
| --- | --- | --- | --- | --- |
| **(Intercept)** | **52.42** | **15.48** | **3.39** | **0.001** |
| *female* vs male | -9.45 | 16.07 | -0.59 | 0.56 |
| enclosure size | -0.14 | 0.11 | -1.26 | 0.21 |
| *neutral* vs mirror | 25.56 | 14.59 | 1.75 | 0.08. |
| ***neutral* vs enrichments** | **31.29** | **12.64** | **2.48** | **0.01** |
| ***neutral* vs predators** | **-67.89** | **12.64** | **-5.37** | **< 0.001** |
| Housing type | 19.10 | 16.73 | 1.14 | 0.25 |
| duration of rehab | 0.62 | 8.05 | 0.08 | 0.94 |

Italic indicates reference level

#### Proximity of individuals to novel objects

The proximity of *N. javanicus* to novel objects varied significantly among individuals (supplemental Table S3). Females (mean ± SD: 0.49 ± 0.25 m) tended to stay closer to novel objects than males (mean ± SD: 1.12 ± 1.41 m), although this effect was not statistically significant. Proximity was strongly influenced by object type: individuals approached the conspecific stimulus (mirror) and enrichment items more closely than the predator models and the neutral object (supplemental Table S3). Housing condition did not have a significant effect on proximity.

Additionally, a Spearman correlation test revealed a significant negative correlation (rho = -0.71; p < 0.01) between boldness score and minimum approach distance to novel objects, with bolder individuals maintaining closer distances to the objects (supplementary Figure S2). However, we need to note that the results did not consider the cage size.

#### Personality and Behavioral Syndromes

We measured the minimum distance each *N. javanicus* approached a novel object and the latency (time to reach) their closest proximity (supplemental Figure S3). Both measures varied widely among individuals and across object types. For example, individual JS04 consistently approached novel objects immediately after noticing them, whereas JS10 and JS05 displayed hesitation before approaching. Other individuals readily approached certain objects but were more cautious toward others. These patterns suggest apparent individual differences in boldness, curiosity, and decision-making strategies when encountering novelty.

Principal component analysis identified two primary behavioral axes (Figure 3). PC1 (eigenvalue = 3.15; 52.2% of variation) represented an activity–boldness gradient, characterized by positive loadings for locomotion (0.53) and boldness (0.44) and negative loadings for resting (−0.52) and grooming (−0.44). PC2 (eigenvalue = 1.33) explained an additional 22.23% of variance and loaded strongly on alert behavior (−0.73) and stereotypy (−0.60), reflecting an anxiety-vigilance dimension orthogonal to exploratory tendency. Together, these components indicated a potential behavioral syndrome in rehabilitating *N. javanicus*, with individuals varying along bold–active and anxiety–vigilant dimensions. Sampling adequacy was moderate (KMO = 0.65), indicating that the data were appropriate for dimension reduction while warranting cautious interpretation of the underlying structure.

**Figure 3.**
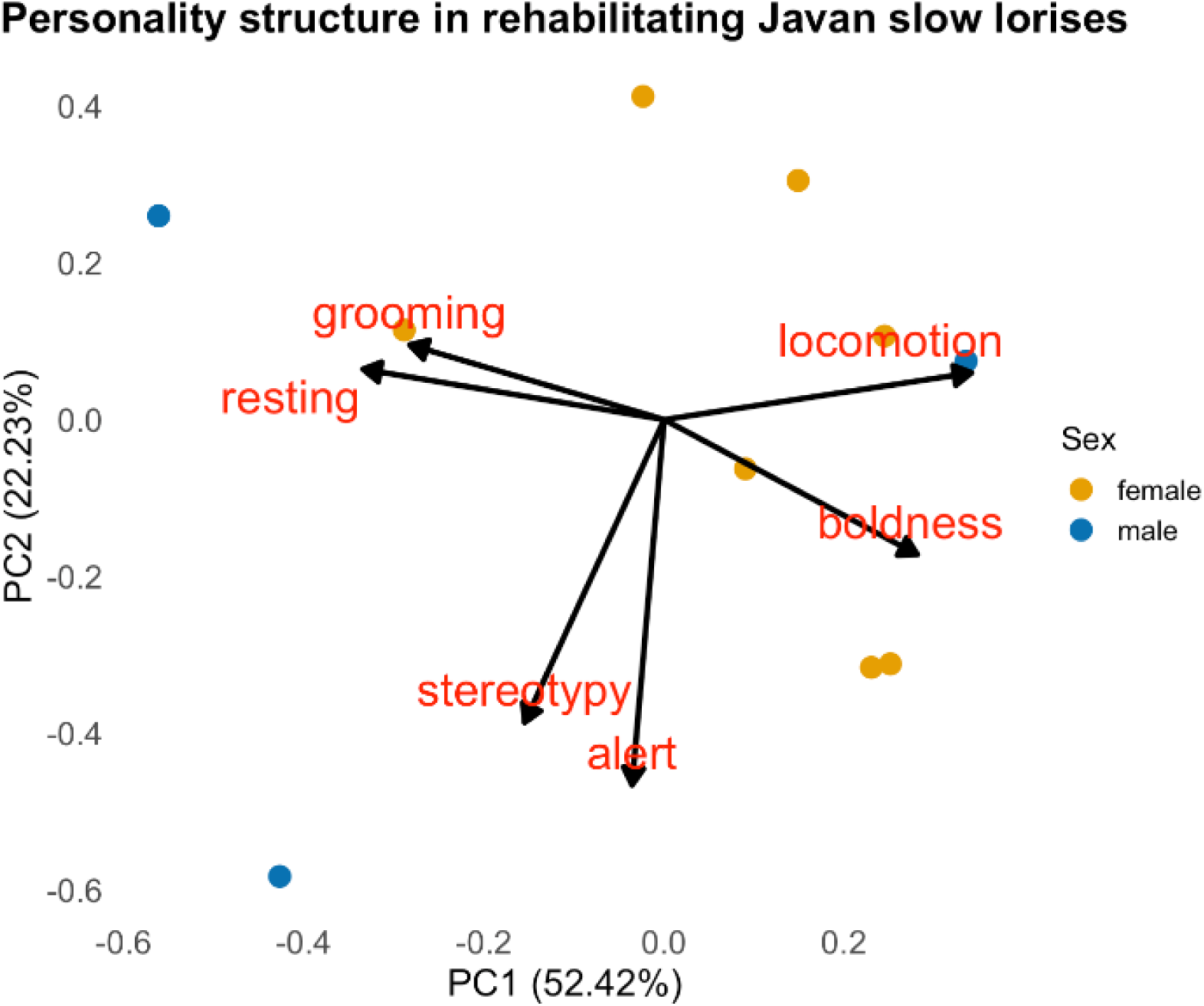
Principal component analysis (PCA) of activity and boldness traits in rehabilitating *N. javanicus*. PC1 indicates a bold–exploratory activity axis, whereas PC2 reflects an anxiety–vigilance axis. Points show individual behavioral profiles.

## Discussion

This study integrates activity budgets and measures of boldness to examine how daily activity patterns and responses to novelty jointly inform behavioral welfare and release readiness in rehabilitating *N. javanicus*. Overall, individuals exhibited locomotion-dominated activity budgets, while responses to novel objects were strongly stimulus-dependent, and multivariate analyses indicated that activity patterns and boldness covaried along an integrated behavioral axis.

### Activity budget

#### Overall

The activity budget of rehabilitating *N. javanicus* was dominated by locomotion (∼57%), consistent with locomotion-dominated budgets reported for *N. javanicus* in both wild and captivity settings (Haris 2008; Priatna et al. 2023; Reinhardt et al. 2016), suggesting that individuals were able to sustain movement patterns, potentially facilitated by the availability of semi-natural climbing structures and enrichment. Rest accounted for the second-largest proportion (∼23%), matching the species’ nocturnal rhythm but appearing higher relative to some wild conspecific reports (Priatna et al. 2023; Reinhardt et al. 2016). Compared with other nocturnal primates such as owl monkeys (*Aotus* spp.), which may devote substantially more time to resting during the night, rehabilitating *N. javanicus* in this study showed a more active nocturnal profile (Link et al. 2023).

Stereotypic movements were relatively infrequent (4.36%), suggesting that current enrichment protocols may mitigate abnormal repetitive behaviors, although not entirely eliminate them. Compared to the wild individuals (3 – 14%) (Reinhardt et al. 2016; Romdhoni et al. 2022), limited feeding time (∼3.13%) at the rehabilitating center likely reflects provisioning schedules rather than natural foraging patterns, a known limitation of captive diets in nocturnal primates (Williams et al. 2015). Social interactions and grooming were observed at low levels (∼2–3%), consistent with slow lorises’ flexible but generally solitary behavior (Nekaris and Bearder 2011; Yamanashi et al. 2021).

We should note, however, that activity and stereotypy patterns in captive or rehabilitative settings are shaped by multiple interacting factors, including enclosure size and complexity, human presence, enrichment type, feeding regimes, social environment, and individual history (Moore et al. 2015). These factors can influence both the expression and timing of behaviors, meaning that direct quantitative comparisons with wild activity budgets should be made cautiously. Nonetheless, the predominance of locomotion across settings suggests that rehabilitated individuals retain core aspects of species-typical activity patterns, even though variation in stereotypy may reflect context-dependent stressors rather than enrichment effects alone (Moore et al. 2015).

#### Sex difference

Females exhibited significantly higher locomotion than males. This pattern is broadly consistent with activity profiles reported for wild *N. javanicus* (Reinhardt et al. 2016), although contrasting evidence exists for Sunda slow lorises (*N. coucang*), where females are more inactive under captive conditions (Ramadhan et al. 2024). Such divergence may reflect species-specific socioecology or variation in rehabilitation environments, and enrichment studies further show that locomotor activity in slow lorises can increase under more complex housing and foraging opportunities (Al-Razi et al. 2020; Williams et al. 2015; Yamanashi et al. 2021). Together, these findings highlight the importance of environmental context in shaping behavioral expression and underscore the need for comparative work across taxa and settings.

#### Housing condition

Although the Dirichlet analysis did not detect a statistically significant effect of housing conditions on overall activity budgets, descriptive patterns suggested differences between solitary and grouped individuals. Solitarily housed *N. javanicus* tended to spend a slightly higher proportion of time resting, whereas grouped individuals showed reduced resting alongside increased social behaviors, indicating a redistribution of activity rather than a uniform suppression of inactivity. Similar patterns have been reported in pygmy slow lorises, where group housing promoted affiliative interactions and greater behavioral diversity following social introductions (Alejandro et al. 2021; Yamanashi et al. 2021). In addition, stereotypic behaviors were descriptively more frequent in solitary individuals, particularly males, suggesting that social isolation may limit behavioral outlets or elevate stress even when differences are not statistically significant (Moore et al. 2015). Previous studies have demonstrated that increased environmental and social complexity, such as group housing, naturalistic climbing structures, and enrichment-based feeding, can reduce abnormal repetitive behaviors and promote more diverse activity patterns in captive *Nycticebus* spp. (Al-Razi et al. 2020; Williams et al. 2015). Taken together, these findings suggest that group housing, especially when combined with appropriate environmental enrichment, may confer welfare benefits by promoting more balanced activity patterns and reducing the expression of stereotypy. Therefore, group housing should be encouraged in rehabilitation settings when individual compatibility and management constraints allow.

### Boldness

The negative correlation observed between boldness scores and proximity to novel objects indicates that bolder individuals tended to investigate unfamiliar stimuli at closer distances, while shy individuals maintained greater separation. This pattern aligns with established bold versus shy behavioral gradients described in other vertebrates, including trout (Frost et al. 2007), vervet monkeys (Blaszczyk 2017), and cross-taxa personality frameworks (Sih et al. 2004b). Behavioral responses were strongly dependent on the type of stimulus presented. Predator models elicited avoidance, whereas enrichment and neutral objects encouraged closer investigation, demonstrating that boldness is expressed conditionally rather than uniformly across contexts. Similar stimulus-dependent responses have been reported in chacma baboons (Carter et al. 2012) and in translocation studies of San Joaquin kit foxes (Bremner-Harrison et al. 2004), supporting the interpretation that boldness is context specific and shaped by perceived risk and reward.

In our analysis, boldness and minimum approach distance were strongly negatively correlated, indicating that bolder individuals tend approached novel stimuli more closely. However, this relationship did not account for enclosure size, and our results suggest that housing space alone is not a significant predictor of boldness. This implies that enclosure size may modulate expression of exploratory behaviour indirectly, rather than driving boldness itself. Larger enclosures may provide greater opportunity for positional avoidance, allowing shy individuals to maintain distance without confronting stimuli. Similar patterns of environment moderated exploration have been reported in slow lorises and other nocturnal primates where enrichment and structural complexity influence activity allocation rather than personality state (Al-Razi et al. 2020; Williams et al. 2015; Yamanashi et al. 2021). In contrast, non-arboreal experimental species such as zebrafish and laboratory mice frequently show increased exploration in larger spaces (An et al. 2021; Maierdiyali et al. 2020), highlighting a potential niche specific divergence in how spatial freedom interacts with risk response. These results underscore the importance of distinguishing personality expression from environmental constraint when evaluating boldness for release planning.

Sex, social housing, and rehabilitation duration did not predict boldness, indicating that observed bold–shy variation in rehabilitating *N. javanicus* was not explained by short-term environmental or demographic factors. However, because each individual was exposed to each stimulus only once, repeatability of boldness could not be formally assessed. Accordingly, the present results should be interpreted as preliminary indicators of personality-related variation, rather than as evidence of stable, repeatable personality traits (Reale et al. 2010). The absence of sex effects contrasts with patterns reported in other mammals where males often exhibit greater risk-taking (Schuett et al. 2010), suggesting either phylogenetic differences or that uniform captive conditions reduce expression of sex-linked behavioral strategies. Similarly, social housing did not increase boldness despite its welfare benefits for *N. pygmaeus* in group settings (Yamanashi et al. 2021), implying that boldness may be less sensitive to social context than activity or affiliative behavior. Taken together, our results support the interpretation that boldness in rehabilitating *N. javanicus* reflects inter-individual differences that are not readily explained by short-term captive conditions.

Latency and minimum approach distance to novel stimuli varied among individuals, suggesting inter-individual differences in responses sto novelty within this study context. Although repeatability was not formally assessed, similar axes of personality variation and repeatable behavioral tendencies have been well documented in primates, including macaques (Adams et al. 2015; Arlet et al. 2024; Neumann et al. 2013) and common marmosets, where exploration and approach latency show consistent individual signatures (Koski and Burkart 2015).

Additionally, the identification of a bold-exploratory behavioral syndrome suggests that activity budgets and boldness responses form integrated personality profiles rather than independent traits (Figure 3). This integration implies consistent behavioral strategies across contexts; individuals that are more active in their enclosures also tend to approach novelty more boldly. Such syndromes have important implications for reintroduction: bold-exploratory individuals may adjust rapidly to unfamiliar environments but incur higher risks, whereas shy-inactive individuals may benefit from caution but require longer acclimation periods (Haage et al. 2017; Sih et al. 2004b). Ecologically, bold-exploratory phenotypes may promote dispersal, foraging innovation, and post-release exploration, while shy-inactive phenotypes may enhance vigilance and predator avoidance (Bremner-Harrison et al. 2004; Cote et al. 2010; Webster and Lefebvre 2001). Recognizing these alternative strategies would inform individualized release planning and improve post-release outcomes in *N. javanicus* rehabilitation.

## Conservation implications

Our findings highlight the potential relevance of personality variation for *N. javanicus* rehabilitation, though the predictive value of these behavioral profiles remains to be validated, and post-release tracking across personality types will be essential to refine and substantiate personality-informed release protocols. Individuals that approach novelty readily may be better equipped to explore unfamiliar landscapes and locate resources post-release, whereas more cautious individuals may benefit from release strategies that minimize predation risk or allow gradual acclimation. Incorporating such behavioral profiles into candidate selection would align with evidence from other primates showing that personality differences predict adaptability and learning capacity (Carter et al. 2013; King and Figueredo 1997; Neumann et al. 2013).

The strong modulation of boldness by stimulus type suggests that pre-release conditioning could be improved by exposing lorises to carefully controlled risk–reward contexts, allowing them to practice discrimination between threats and neutral or beneficial stimuli. Targeted enrichment, including ecologically meaningful foraging challenges, has been shown to shape engagement and reduce abnormal behavior in nocturnal primates (Al-Razi et al. 2020; Williams et al. 2015; Yamanashi et al. 2021). Accordingly, any conditioning protocols involving risk-related stimuli should be closely monitored and adapted to minimize distress.

Looking forward, integrating pre-release personality metrics with post-release monitoring would allow direct evaluation of whether boldness and activity syndromes predict exploration, resource acquisition, or survival after release. A personality-informed framework could support adaptive release matching, whereby bolder-explorative individuals are released into resource-rich habitats with lower immediate predation risk or greater canopy connectivity, while shyer-inactive individuals are matched to structurally complex forests that facilitate refuge use and cautious dispersal. In rescue–release scenarios where extended rehabilitation is not feasible, this same framework could be implemented through brief behavioral screening using standardized novel-object or emergence tests may provide a practical means of informing release decisions while minimizing handling time and stress.

## Acknowledgements

We would like to thank Kyoto University Wildlife Research Center (WRC), CICASP, Research Units for Exploring Future Horizons (Coevolution and Coexistence), and the Joint Research Program of WRC for facilitating our research, YIARI Bogor board of directors and staff (especially Mastur, Ajo, Aconk, Ganyong, Pak Aki, Kojek) for their permission and assistance in the field, FMIPA-Biologi IPB University dean and staff. We also thanked BRIN, KLHK, and BBKSDAE Jawa Barat for the administrative assistance and permission to conduct research in Indonesia.

## Author contributions

AL: Conceptualization, formal analysis, methodology, investigation, writing – original draft, funding acquisition, project administration. WP: writing – review & editing, project administration, investigation. NPP: writing – review & editing, project administration, investigation. PR: writing – review & editing, project administration. RM: writing – review & editing, project administration. MS: conceptualization, writing – review & editing, methodology, supervision, funding acquisition. IM: formal analysis, writing – original draft, supervision, funding acquisition. AM: conceptualization, writing – review & editing, methodology, supervision, funding acquisition.

## Data availability

All data needed to evaluate the conclusions in the paper are present in the paper. Additional data related to this paper may be requested from the authors.

## Declarations

### Ethical approval

This research was conducted in accordance with the Guidelines for the Care and Use of Non-human Primates and the Guidelines for Field Research on Non-human Primates established by the Center for the Evolutionary Origins of Human Behavior, Kyoto University. Ethical approval and permissions were obtained from the Field Research Committee of the Wildlife Research Center, Kyoto University, as well as the Indonesian National Research and Innovation Agency (BRIN, Ref No.: 009/KE.02/SK/01/2024)).

### Generative AI and AI-assisted technologies in the writing process

During the preparation of this manuscript, the authors used ChatGPT-5.1 to refine the clarity and logical flow of the text. The authors carefully reviewed, corrected, and approved all content generated, and take full responsibility for the final published version.

