## Supplemental Table S1 for "Activity Budgets and Boldness as Potential Predictors of Release Readiness in Rehabilitating Javan Slow Lorises"

*correspondence to: abdullahlanggeng.gmail.com

**SUPPLEMENTARY**

Supplemental Table S1. Housing and demographic attributes of rehabilitating *N. javanicus*. Sex, group/solitary condition, enclosure size, rehabilitation duration and total observation time are shown for each subject.

| No | ID | Sex | Origin | Date of arrival in rescue center (DD/MM/YY) | Enclosure Size  (m^2^) | Housing type | Conspecific | Total observation minutes |
| --- | --- | --- | --- | --- | --- | --- | --- | --- |
| 1 | JS01 | Female | BKSDA Yogyakarta | 26/01/22 | 16 | Solitary | - | 540 |
| 2 | JS02 | Female | BKSDA Yogyakarta | 15/01/24 | 36 | Group | JS08 | 360 |
| 3 | JS03 | Female | BKSDA Ciamis | 20/10/23 | 16 | Solitary | - | 540 |
| 4 | JS04 | Male | BKSDA Yogyakarta | 15/01/24 | 16 | Solitary | - | 405 |
| 5 | JS05 | Female | Jonggol | 17/07/22 | 192 | Group | JS10 | 495 |
| 6 | JS06 | Female | BKSDA Yogyakarta | 15/01/24 | 36 | Group | Non-releasable individual | 360 |
| 7 | JS07 | Female | BKSDA Bogor | 20/12/23 | 192 | Solitary | - | 495 |
| 8 | JS08 | Female | BKSDA Soreang | 15/01/24 | 36 | Group | JS02 | 360 |
| 9 | JS09 | Male | BKSDA Yogyakarta | 15/01/24 | 16 | Solitary | - | 495 |
| 10 | JS10 | Male | TNGHS Kabandungan | 02/09/22 | 192 | Group | JS05 | 495 |

Supplemental Table S2. Summary of boldness score for individuals exposed to each novel object and closest proximity to the object.

| ID | Sex | Boldness score and closest proximity (m) | | | | | | | | | | | | Mean boldness score | Mean Proximity (m) |
| --- | --- | --- | --- | --- | --- | --- | --- | --- | --- | --- | --- | --- | --- | --- | --- |
|  |  | Snake | | Leopard Cat | | Enrichment 1 | | Enrichment 2 | | Pyramids | | Mirror | |  |  |
|  |  | Score | Proximity | Score | Proximity | Score | Proximity | Score | Proximity | Score | Proximity | Score | Proximity |  |  |
| JS01 | Female | 0.9 | 0.8 | -18.7 | 0.7 | 54.8 | 0 | 22.9 | 0 | 86.1 | 0.8 | 103.7 | 0 | 49.6 | 0.3 |
| JS02 | Female | -58.7 | 0.6 | 3.8 | 0.5 | 76.8 | 0 | 138.4 | 0 | 56.6 | 0.7 | 84.6 | 0 | 54.4 | 0.3 |
| JS03 | Female | -62.6 | 0.6 | -48.9 | 1.0 | 72.6 | 0 | 70.6 | 0 | 77.1 | 0.5 | 90.3 | 0 | 42.3 | 0.3 |
| JS04 | Male | 99.1 | 0.3 | 12.9 | 0.2 | 109.4 | 0 | 111.6 | 0 | 72.6 | 0.0 | 41.3 | 0 | 71.7 | 0.1 |
| JS05 | Female | -7.0 | 0.9 | 55.0 | 0.7 | 156.0 | 0 | 113.1 | 0 | 46.8 | 0 | 102.0 | 0.3 | 69.2 | 0.6 |
| JS06 | Female | -76.7 | 1.9 | -37.1 | 1.6 | 48.9 | 0 | 39.1 | 0 | 60.1 | 1.2 | 55.3 | 0 | 27.9 | 0.7 |
| JS07 | Female | -66.2 | 1.4 | -55.8 | 2.3 | 67.4 | 0.3 | 26.7 | 1.1 | 52.6 | 0.3 | 81.6 | 0 | 15.0 | 0.9 |
| JS08 | Female | -1.9 | 0.3 | 9.9 | 0.3 | 79.6 | 0 | 140.9 | 0 | 51.6 | 1.4 | 95.6 | 0 | 61.8 | 0.3 |
| JS09 | Male | -57.4 | 1.1 | -43.9 | 0.3 | 96.0 | 0 | 112.1 | 0 | -24.7 | 1.4 | 81.6 | 0 | 14.6 | 0.5 |
| JS10 | Male | -20.2 | 5.8 | -40.9 | 3.8 | 18.1 | 0 | 14.7 | 0 | -7.0 | 2.4 | -8.4 | 2.1 | -12.3 | 2.7 |
| Mean per novel object | | -25.1 | 1.4 | -16.4 | 1.2 | 78.0 | 0.1 | 79.0 | 0.1 | 47.2 | 0.9 | 72.8 | 0.2 | - | - |

Supplemental Table S3. Generalized Linear Mixed Model (GLMM) results for minimum approach distance to novel objects. Predictors included object type, sex, housing type, and duration in rehabilitation, with individual identity and sampling date as random effects. Significant predictors indicated in bold (p ≤ 0.05).

|  | Estimate | Std. error | z value | Pr(>\|z\|) |
| --- | --- | --- | --- | --- |
| (Intercept) | 0.59 | 0.32 | 1.85 | 0.07. |
| *female* vs male | 0.57 | 0.37 | 1.53 | 0.13 |
| ***neutral* vs mirror** | **-0.64** | **0.29** | **-2.19** | **0.03** |
| ***neutral* vs enrichments** | **-0.78** | **0.25** | **-3.07** | **0.002** |
| *neutral* vs predators | 0.37 | 0.25 | 1.47 | 0.14 |
| Housing type | 0.29 | 0.36 | 0.80 | 0.42 |
| duration of rehab | 0.20 | 0.18 | 1.13 | 0.26 |

Italic indicates reference level

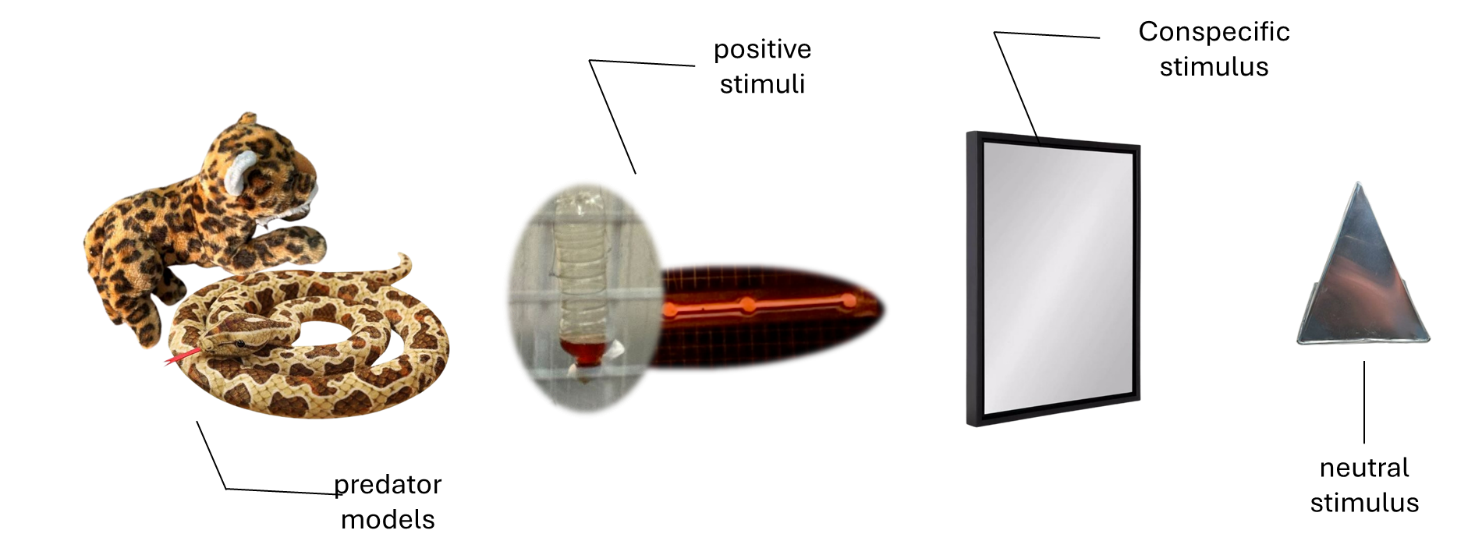

Supplemental Figure S1. Novel objects used to assess boldness in rehabilitating *N. javanicus*. Objects represented four stimulus categories: predator cues (snake model; leopard cat plush), enrichment items (honey bottle; bamboo fruit puzzle), conspecific stimulus (mirror), and neutral object (pyramid). Each individual was exposed once to each object under standardized conditions.

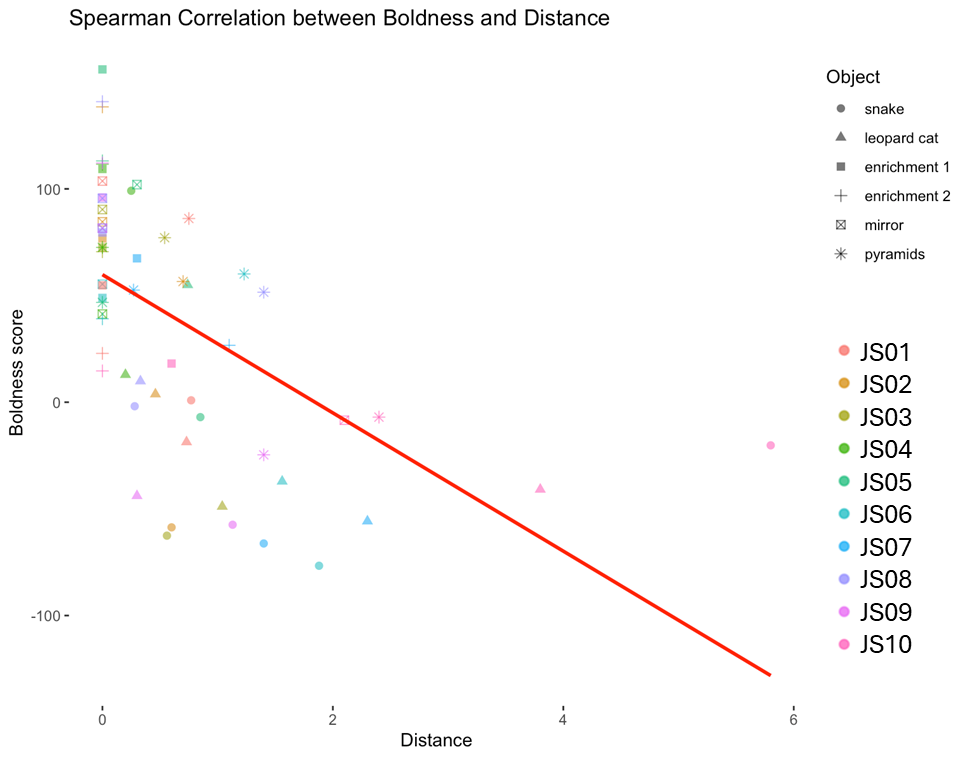

Supplemental Figure S2. Relationship between boldness score and minimum approach distance. Each point represents the mean response of one individual across stimuli. A significant negative correlation indicates that bolder *N. javanicus* approached objects more closely (Spearman’s rho = −0.71, p < 0.001).

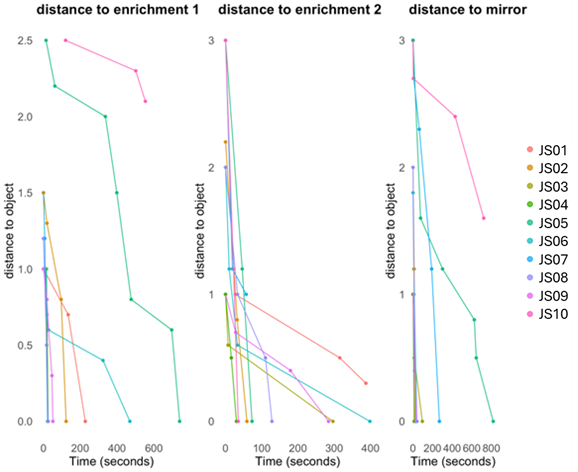

Supplemental Figure S3. Individual variation in approach latency and minimum distance to novel objects. Lines indicate per-individual response to three different objects. Shorter latencies and distances denote bolder behavioral tendencies.
